# Long-term SARS-CoV-2 Persistence in Syrian Hamsters

**DOI:** 10.64898/2026.07.19.739454

**Authors:** Thais Melquiades de Lima, Carlos Eduardo Capelini Eli Lopes, Maria Vitoria Oliveira de Souza, Felipe Rocha do Nascimento, Daniela Méria Ramos Rodrigues, Gabriela Conde Silva, Matheus Dias, Tufi Antônio Nasser Neto, Maria Lúcia Silva, Daniel Macedo de Melo Jorge, Juliano de Paula Souza, Eurico Arruda

## Abstract

SARS-CoV-2 persistence has been proposed as a potential contributor to the pathogenesis of long COVID, with reservoir tissues potentially serving as sites for viral persistence, intra-host evolution, and intermittent viral shedding. Here, we used experimentally infected Syrian hamsters to investigate long-term SARS-CoV-2 persistence across tissues, viral infectivity, and associated immunological and metabolic alterations. Syrian hamsters (Mesocricetus auratus) were intranasally infected with a SARS-CoV-2 parental strain or Gamma and Delta variants and monitored for up to one year, with samples collected at 3, 15, 30, 90, 150, and 365 days post-infection (dpi). During the acute phase, infected animals exhibited significant weight loss, viral shedding, and marked pulmonary inflammation, accompanied by increased expression of pro-inflammatory cytokines at 3 dpi. Infection was confirmed by seroconversion, with sustained IgG responses and low-titer neutralizing antibodies against Omicron. Viral nucleoprotein was detected in multiple tissues up to 365 dpi, while RT-qPCR identified persistent low-level viral RNA in the lungs, brain, spleen, and thymus throughout the observation period, without evidence of productive viral replication. Immune gene expression displayed organ-specific temporal patterns: acute pulmonary inflammation transitioned into broad late-stage suppression, except for sustained TGF-β expression; the brain exhibited a late chemokine signature at 365 dpi; and the thymus showed a delayed immune activation peak at 150 dpi, particularly in Delta-infected animals. Metabolomic profiling revealed a shared acute-phase metabolic signature across variants that largely resolved by 365 dpi, whereas Delta-infected animals retained distinct residual metabolic alterations. Collectively, these findings establish a model of long-term SARS-CoV-2 tissue persistence characterized by organ-specific immune and metabolic signatures, providing a platform to investigate mechanisms underlying post-acute sequelae and evaluate potential therapeutic strategies.

## INTRODUCTION

COVID-19 is primarily a respiratory disease (Guan et al., 2020), yet SARS-CoV-2 infection frequently presents with extra-respiratory manifestations, reflecting its systemic nature and association with multi-organ involvement in severe cases (Liu et al., 2021).

Increasing evidence suggests that the persistence of SARS-CoV-2 in tissues beyond the acute phase of infection may contribute to the development of post-acute sequelae of COVID-19 (long COVID). Prolonged detection of viral RNA and proteins has been associated with sustained immune activation and immune dysregulation, which may contribute to chronic inflammation and long-lasting clinical manifestations (Yang et al., 2023). The presence of viral components months to more than one year after acute infection in individuals with post-acute sequelae of COVID-19 supports the hypothesis that viral persistence may disrupt local immune homeostasis and contribute to persistent symptoms. Collectively, these findings suggest a potential association between tissue persistence of SARS-CoV-2 and an increased risk of long COVID (Craddock et al., 2023; Yang et al., 2023; Zuo et al., 2024).

Golden Syrian hamsters (*Mesocricetus auratus*) are widely used as an animal model for COVID-19 due to their susceptibility to SARS-CoV-2 infection, exhibiting efficient viral replication, viral shedding, weight loss, pulmonary histopathological changes, and robust immune responses that resemble human COVID-19 disease (Rosenke et al., 2020; Sia et al., 2020). In this context, we employed the Syrian hamster model to investigate long-term SARS-CoV-2 tissue persistence and its associated immunological features. We assessed viral RNA loads, the presence of infectious particles and viral components across multiple tissues, as well as humoral immune responses. Despite inherent limitations, the Syrian hamster remains a valuable model for studying SARS-CoV-2 pathogenesis, viral persistence, and mechanisms potentially underlying long COVID.

## MATERIALS AND METHODS

### Viruses

The ancestral SARS-CoV-2 strain B.1.1.33 (GenBank accession SAMN32093250, ID: 32093250) was isolated and sequenced at the Viral Pathogenesis Laboratory, Center for Research in Virology, University of São Paulo, Ribeirão Preto, Brazil. The SARS-CoV-2 variants Gamma P.1 (GenBank accession SAMN32093252, ID: 32093252) and Delta AY.99.2 (GenBank accession SAMN32093254, ID: 32093254) were kindly provided by Prof. Edison Luiz Durigon (Institute of Biomedical Sciences, University of São Paulo, São Paulo, Brazil). Viral stocks were propagated in Vero E6 cells. Virus titers were determined by the 50% tissue culture infectious dose (TCID₅₀) assay in Vero E6 cells incubated at 37°C with 5% CO₂. All experiments involving SARS-CoV-2 were conducted in a biosafety level 3 (BSL-3) laboratory at the Center for Research in Virology, University of São Paulo, Ribeirão Preto, Brazil.

### Experimental infection of syrian hamsters

Male Syrian golden hamsters (*Mesocricetus auratus*), 8–10 weeks old, were obtained from Anilab (Paulínia, São Paulo, Brazil) and housed under BSL-3 conditions for one week prior to the experiments for acclimatization. A total of 96 hamsters were randomly assigned to four groups of 24 animals each: two groups infected with distinct SARS-CoV-2 variants (Gamma and Delta), one group infected with the parental SARS-CoV-2 strain, and one mock-infected control group. Four animals from each group were euthanized at 3, 15, 30, 90, 150, and 365 days post-infection (dpi).

Animals were housed in individually ventilated cages under standardized BSL-3 conditions at the Center for Research in Virology, Ribeirão Preto, Brazil, where all procedures were conducted, with food and water provided *ad libitum*. All experimental protocols involving virus handling and animal infections were approved by the institutional animal ethics and biosafety committees and were conducted in accordance with the guidelines of the Animal Ethics Committee of the Ribeirão Preto Medical School, University of São Paulo (FMRP/USP), under approved protocol number 1045/2022R2.

Hamster infections were performed as previously described (Chan et al., 2020; Sia et al., 2020). Animals were anesthetized with 1.5% isoflurane and intranasally inoculated with 10⁶ TCID₅₀ of SARS-CoV-2 in a total volume of 100 μL, and mock-infected animals received 100 μL of DMEM. Following infection, animals were monitored daily, and body weight was recorded for 10 days post-infection. At the indicated time points, blood serum and tissues including lungs, spleen, thymus, cervical lymph nodes, inguinal lymph nodes, brain, and intestine were collected. Serum samples were stored at −80°C until further use. Brain and lung tissues were divided into four portions: one preserved in TRIzol reagent (Thermo Fisher Scientific) for RNA extraction, one stored in RNAlater® solution (Invitrogen, Carlsbad, CA, USA), one fixed overnight at 4°C in Carnoy’s solution (60% ethanol, 30% acetic acid, and 10% chloroform; Merck, Darmstadt, Germany) for subsequent immunohistochemical analysis, and one frozen in viral transport medium (RPMI supplemented with 20% fetal bovine serum and 15% glycerol) for virus isolation and titration attempts. Thymus, spleen and cervical and inguinal lymph nodes were divided into three portions for TRIzol maceration, RNAlater® preservation, and Carnoy fixation.

### Viral load

Total RNA was isolated from lung, spleen, thymus, and brain samples using TRIzol reagent (Thermo Fisher Scientific), following the manufacturer’s instructions. RNA concentration and purity were assessed using a NanoDrop spectrophotometer (Thermo Fisher Scientific). Subsequently, 1 µg of total RNA was reverse-transcribed into complementary DNA (cDNA) using the High-Capacity cDNA Reverse Transcription Kit (Applied Biosystems™), according to the manufacturer’s protocol.

Detection and quantification of SARS-CoV-2 genome were performed using primer and probe sets targeting the 2019-nCoV N2 gene, the E gene, and the subgenomic E RNA, following protocols established by the U.S. Centers for Disease Control and Prevention (Lu et al., 2020; Immergluck et al., 2021) and the Charité group (Corman et al., 2020). The N2 and E genes were analyzed by real-time RT-PCR using a qPCRBIO Probe Mix (PCR Biosystems) and cDNA as template, while subgenomic E RNA was analyzed using total RNA with the qPCRBIO One-Step Go RT-PCR Kit (PCR Biosystems).

Real-time PCR assays were carried out on a QuantStudio™ 5 Real-Time PCR System (Applied Biosystems, USA). Amplification reactions were performed using specific primers (10 µM), probes (5 µM), and qPCRBIO Probe Mix or qPCRBIO One-Step Go (PCR Biosystems), under the following cycling conditions: 50°C for 10 min and 95°C for 2 min, followed by 40 cycles of 95°C for 5 s and 60°C for 30 s.

The primer and probe sequences used were as follows: N2 forward: 5′-TTACAAACATTGGCCGCAAA-3′; N2 reverse: 5′-GCGCGACATTCCGAAGAA-3′; N2 probe: 5′-FAM-ACAATTTGCCCCCAGCGCTTCAG-BHQ1-3′. E gene forward: 5′-ACAGGTACGTTAATAGTTAATAGCGT-3′; E gene reverse: 5′-ATATTGCAGCAGTACGCACACA-3′; E gene probe: 5′-FAM- ACACTAGCCATCCTTACTGCGCTTCG-BHQ1-3′ Subgenomic E forward: 5′-CGATCTCTTGTAGATCTGTTCTC-3′; Subgenomic E reverse: 5′-ATATTGCAGCAGTACGCACACA-3′; Subgenomic E probe: 5′-FAM-ACACTAGCCATCCTTACTGCGCTTCG-BHQ1-3′.

Viral RNA copy numbers were determined using cDNA generated from viral stocks quantified in TCID5 (50% tissue culture infectious dose) as quantification standards, from which standard curves were generated using serial tenfold dilutions. The N2 gene was selected for viral load quantification due to its higher sensitivity (Figure 1). Ct values above 36 were considered indicative of low-level viral genome and were included in the analysis when consistently detected across technical replicates and in the absence of amplification in negative controls.

**Figure 1.**
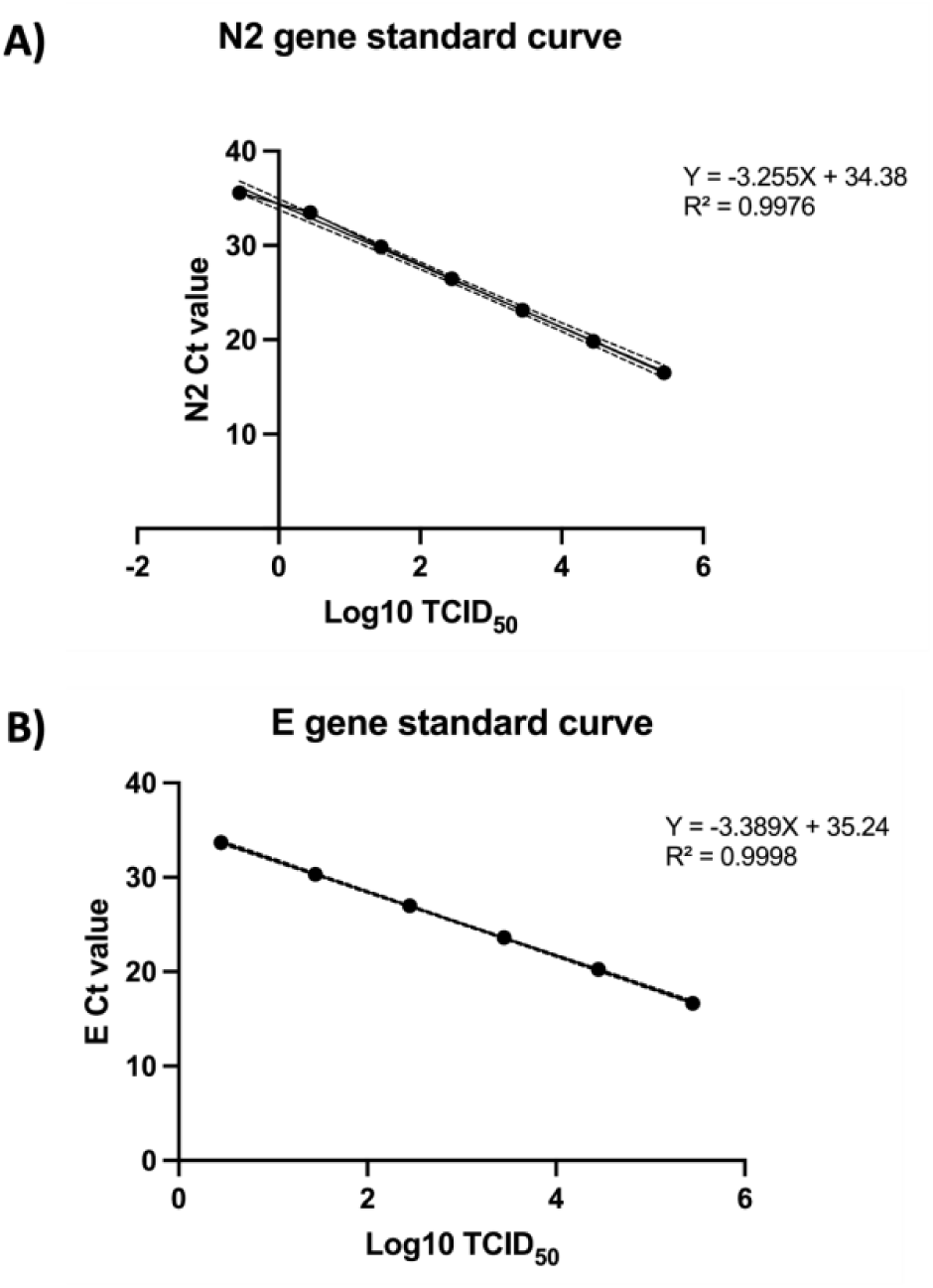
Standard curves for qPCR-based quantification of viral titer. Linear regression curves correlating RT-qPCR Ct values with infectious viral titer (Log10 TCID₅₀/mL), determined from serial dilutions of viral stock with known titer. (A) Standard curve for the N2 gene, with regression equation Y = −3.255X + 34.38 (R² = 0.9976); dashed lines represent the 95% confidence interval of the regression. (B) Standard curve for the E gene, with regression equation Y = −3.389X + 35.24 (R² = 0.9998). Both curves were used to convert Ct values obtained in experimental samples into equivalent viral titers (Log10 TCID₅₀), enabling relative quantification of viral load across the evaluated tissues.

### qPCR for cytokine, inflammatory, and antiviral gene expression

Diluted cDNA samples (1:10) were used for quantitative PCR analysis using the qPCRBIO SybrGreen Mix on a QuantStudio™ 5 Real-Time PCR System (Applied Biosystems). Relative expression levels were expressed as fold change using the formula 2^(-ΔΔCt). Primer sequences used in this study are listed in Supplementary Information (Supplementary Table 1).

### Immunofluorescence on paraffin-embedded tissues

Tissue sections were deparaffinized at 60 °C for 40 min and sequentially rehydrated through xylene, xylene/ethanol, graded ethanol (100%, 95%, 70%, 50%), and rinsed twice in distilled water. Antigen retrieval was performed in Tris-EDTA buffer (10 mM Tris, 1 mM EDTA, 0.05% Tween 20, pH 9.0) using sequential microwave heating, followed by cooling in buffer and washed with PBS washes. Sections were circumscribed with a hydrophobic pen, permeabilized in PBS for 5 min and PBST (0.3% Triton X-100) for 15 min, then washed by 3 × 5 min in PBS. Positive and negative controls were included in this moment (Figure 2). Blocking was performed with 3% goat serum in SuperBlock (Thermo Scientific, Cat No 37515) for 1 h. The primary antibody Rabbit monoclonal [EPR24334-118] to SARS-CoV-2 nucleocapsid protein (1:1000, Abcam, Cat No ab271180) was applied overnight at 4 °C in a humid chamber. The secondary antibody Alexa Fluor 594 goat anti-rabbit IgG (H+L) (1:700, Invitrogen, Cat No A11012), with DAPI (1:1000, Invitrogen, Cat No D21490) and incubated for 1 h at room temperature in the dark, followed by 3 × 5 min PBST washes. Autofluorescence was reduced with 0.1% Sudan Black in 70% ethanol for 15 min in the dark, then washed 3 × 5 min in PBS. Slides were mounted with Fluoromount-G (Invitrogen) and imaged on a Stellaris 8 confocal microscope (Leica Microsystems) at 40× magnification. Images were analyzed using ImageJ.

**Figure 2.**
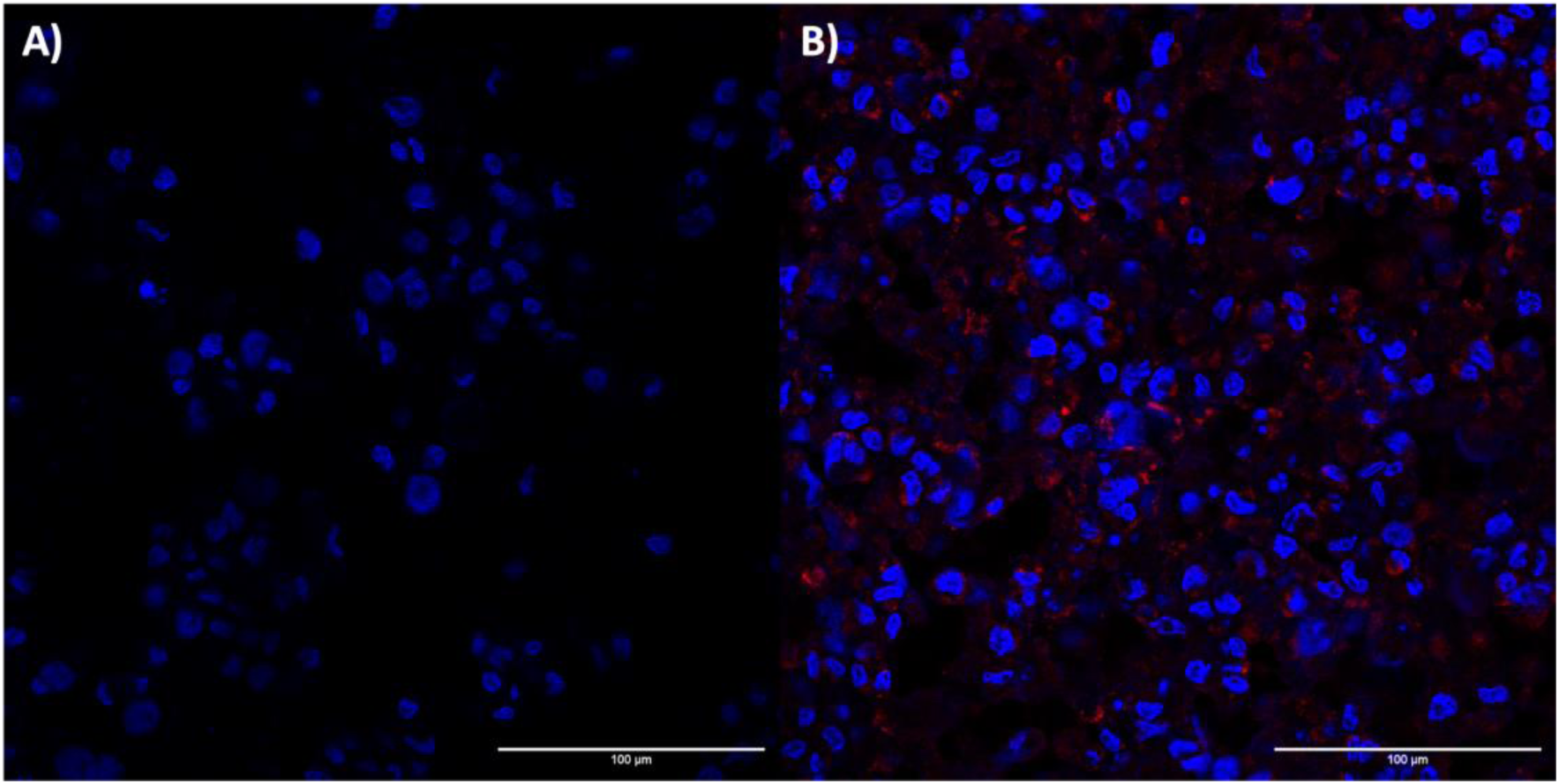
Controls for standardizing immunofluorescence protocol for SARS-CoV-2 NP protein. **A)** Non-infected Vero E6 cells served as a negative control for anti-NP staining, displaying no detectable signal. **B)** SARS-CoV-2-infected Vero E6 cells exhibited a pronounced reddish signal when stained with anti-NP, indicating abundant presence of the NP protein. (magnification of 100 μm).

### Viral isolation samples

Macerated and clarified tissues were stored at −80 °C. Briefly, 100 μL of each clarified sample testing positive for SARS-CoV-2 by RT-qPCR was added to VeroE6/TMPRSS2 cells seeded the day before in a 24-well plate and incubated at 37 °C for 5 days. Cytopathic effects (CPE) were monitored by light microscopy, and viral isolation was considered negative if no CPE was observed after 5 days. Three blind passages were performed for each sample, and the presence of the virus was subsequently confirmed by indirect immunofluorescence. All viral isolation procedures were conducted in a biosafety level 3 (BSL-3) laboratory.

### Histological analysis

Paraffin-embedded tissue blocks were sectioned at 3 μm and mounted on charged glass slides. Sections were deparaffinized in xylene and rehydrated through a graded ethanol series. For hematoxylin and eosin (H&E) staining, slides were immersed in 0.1% Harris’s hematoxylin (Sigma-Aldrich, Cat No 1092531000) for 10 min, followed by 0.5% eosin Y (Sigma-Aldrich, Cat No E4009). Slides were then dehydrated through ascending ethanol concentrations (50%, 70%, 95%, 100%), cleared in xylene, and coverslipped by Entellan new (Merck Millipore, Cat No 1079610100). Tissue morphology and pathological features were evaluated by a board-certified pathologist.

### ELISA assays

Anti–SARS-CoV-2 IgG antibodies present in hamster serum were quantified using an indirect ELISA. For this analysis, the Hamster Anti–SARS-CoV-2 (Receptor Binding Domain, RBD) IgG ELISA Kit (FineTest, Cat. No. EHA0026) was used according to the manufacturer’s instructions. Serum samples were diluted 1:100 prior to analysis. IgG concentrations were calculated based on the standard curve and corrected for the dilution factor. The absorbance was measured at 450 nm using a Varioskan LUX microplate reader (Thermo Fisher Scientific). Data were analyzed using SkanIt software (Thermo Fisher Scientific) and Microsoft Excel (Microsoft).

### Viral microneutralization assay

To evaluate antibody neutralization against the SARS-CoV-2 Omicron variant, serum samples collected from hamsters at different time points post-infection were analyzed using a microneutralization assay. The assay was performed in Vero ACE2/TMPRSS2 cells. Prior to testing, serum samples were heat-inactivated at 56 °C for 30 min. Briefly, 50 μL of twofold serial dilutions of each serum sample (starting at 1:10) were mixed with 50 μL containing 100 TCID₅₀ of SARS-CoV-2 and incubated at 37 °C for 1 h to allow virus neutralization. The serum–virus mixtures were then added to confluent Vero ACE2/TMPRSS2 cell monolayers and incubated at 37 °C with 5% CO₂ for 96 h. After incubation, cells were fixed with 10% formaldehyde for 20 min and stained with crystal violet for visualization of cytopathic effects (CPE). Neutralizing antibody titers were determined as the highest serum dilution that completely inhibited CPE in duplicate wells. A serum dilution ≥1:10 was considered positive for neutralizing activity. Assay validation included a serum sample previously confirmed to neutralize the Omicron variant as a positive control and a pre-pandemic serum sample as a negative control (Campos et al., 2022; Jeong et al., 2024).

### Metabolomics analysis

Hamster serum samples, stored at −80°C, were extracted using acetonitrile and chilled methanol following the protocol established by the Mass Spectrometry and Organic Molecules Center at the School of Pharmaceutical Sciences. Briefly, serum was solubilized in a chilled extraction solution (methanol:acetonitrile, 1:1), vortexed for one minute, stored at −20°C for 30 minutes, and centrifuged at 20,000 × g at 4°C for 10 minutes. The supernatant was then collected, evaporated to dryness at 25°C in a vacuum concentrator, and resuspended in the initial mobile phase ratio prior to injection into a timsTOF fleX MALDI-2 mass spectrometer (Bruker) with ion mobility acquisition. Raw data in .d format were imported into MZmine 3 (Schmid et al., 2023) and processed according to the software’s standard workflow for LC-IMS-MS data.

Import included software-based centroiding, and mass detection was performed using the “lowest signal factor” method, with noise thresholds of 500 for MS1 and 100 for MS2, determined after visual inspection. Chromatograms were constructed using the Modular ADAP Chromatogram Builder algorithm. Following complete processing, chromatograms were spectrally aligned against the FragHub library (Dablanc et al., 2024) within MZmine 3, retaining hits with a cosine score greater than 0.7. Unidentified chromatograms were submitted to SIRIUS (Dührkop et al., 2019) using default matching and deduction parameters to maximize metabolite annotation, and all hits with a score greater than 0.75 were retained following manual curation.

### Statistical analysis

Statistical analyses were performed using GraphPad Prism (version 10.2.1 (339), GraphPad Software, San Diego, CA, USA). Percentage body weight change, viral load (N2 gene copy number), and mRNA expression levels were analyzed using two-way analysis of variance (Two-Way ANOVA), or mixed-effects analysis when missing data were present, with viral variant and time post-infection as factors, followed by Dunnett’s test to compare each infected group with mock at each time point. For mRNA expression, fold-change values (2^−ΔΔCt) were log2-transformed prior to analysis, and temporal kinetics within each variant were additionally assessed using Tukey’s test. Differences were considered statistically significant when p < 0.05 (*p < 0.05, **p < 0.01, ***p < 0.001, ****p < 0.0001).

## RESULTS

### Body weight changes following infection with SARS-CoV-2 variants

To assess disease progression following SARS-CoV-2 infection, body weight was recorded daily for 10 days in hamsters assigned to four experimental groups: Mock (n = 24), Parental strain (n = 24), Gamma variant (n = 24), and Delta variant (n = 24). Four animals per group were euthanized at each scheduled time point for tissue collection and subsequent analyses.

Infection resulted in variant-dependent changes in body weight (Figure 3). Hamsters infected with the Parental strain or the Gamma variant experienced greater and more sustained weight loss than those infected with the Delta variant, which displayed only a modest and transient decrease in body weight, followed by recovery by day 10 post-infection. As expected, Mock animals showed no significant changes in body weight throughout the study period.

**Figure 3.**
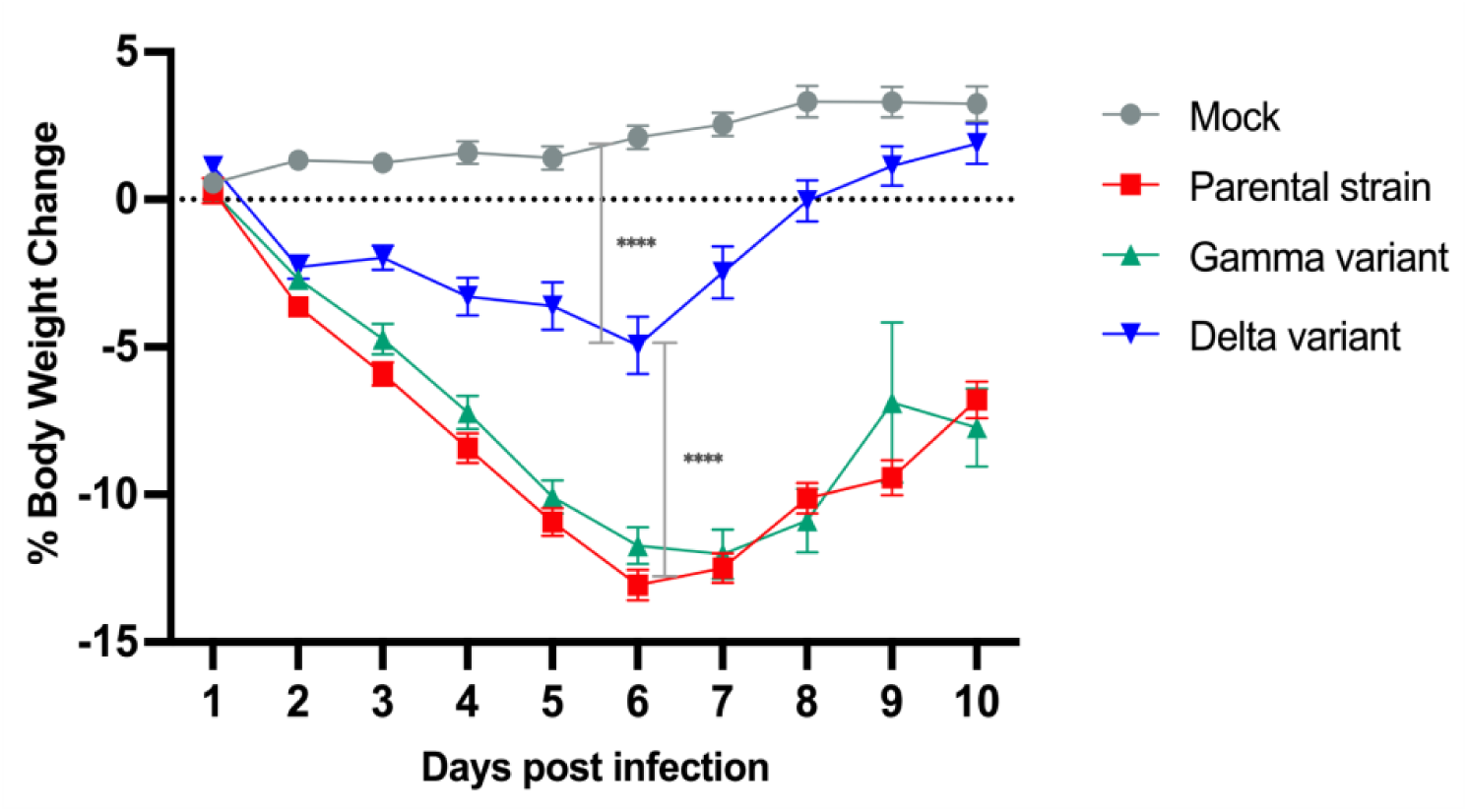
Body weight changes following SARS-CoV-2 infection in Syrian hamsters. Percentage of body weight change in mock-infected and SARS-CoV-2-infected hamsters inoculated with the parental strain, Gamma, or Delta variant during the first 10 days post-infection (dpi). Data are presented as mean ± SEM. Statistical significance was determined by two-way ANOVA with Tukey’s multiple comparisons test. \*\*\*\**P* < 0.0001 for Delta variant versus mock and versus the parental strain and Gamma variant.

### The lung exhibits the highest viral burden and persistent viral RNA detection

To investigate viral dissemination and persistence, viral RNA was quantified by RT-qPCR targeting the SARS-CoV-2 N2 gene in the lungs, brain, spleen, and thymus collected at 3, 15, 30, 90, 150, and 365 dpi. Standard curves demonstrated high amplification efficiency and linearity for both the N2 and E assays (Figure 1). The N2 assay showed greater analytical sensitivity, detecting viral RNA down to 0.2 TCID50, and was therefore used for viral load quantification. Viral RNA was detected in all tissues during the acute phase and remained detectable at low levels in a subset of animals at later time points consistent with low-level viral persistence.

The lung exhibited the highest viral RNA levels during the acute phase of infection. At 3 dpi, all infected groups (Parental, Gamma, and Delta) showed high viral RNA levels, ranging from approximately 10⁵ to 10⁶ N2 copies/μL cDNA (Figure 4A). Viral RNA levels appeared to decline progressively over time, reaching lower levels between 90 and 150 dpi. Nevertheless, low levels of viral RNA (approximately 10–50 N2 copies/μL cDNA) remained detectable in a subset of animals at 365 dpi, indicating the persistence of viral RNA up to one year after infection.

**Figure 4.**
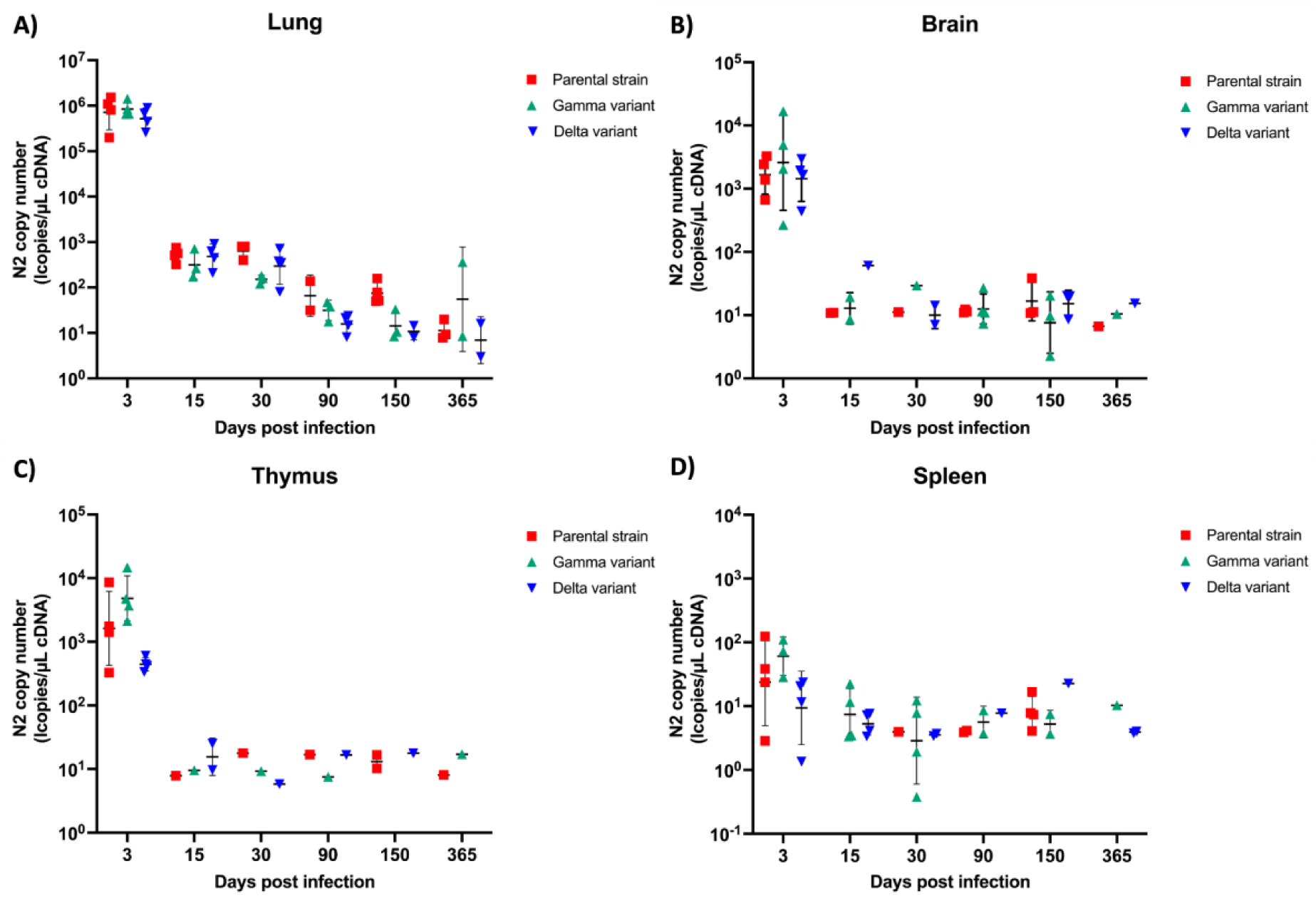
Long-term persistence of SARS-CoV-2 RNA in tissues of infected Syrian hamsters. SARS-CoV-2 N2 RNA copy numbers determined by RT-qPCR in the **A)** lung, **B)** brain, **C)** thymus, and **D)** spleen at 3, 15, 30, 90, 150, and 365 days post-infection (dpi). Hamsters were infected with the parental strain, Gamma, or Delta variant. Each symbol represents an individual animal, and horizontal bars indicate the mean ± SD. Viral RNA levels are expressed as copies/μL cDNA and plotted on a logarithmic scale.

### Distinct viral RNA kinetics in extrapulmonary tissues with persistent viral detection

Extrapulmonary tissues also harbored viral RNA throughout the study, although at lower levels than those observed in the lungs.

In the brain, viral RNA was readily detected at 3 dpi, with levels declining markedly after 15 dpi to approximately 10–20 N2 copies/μL cDNA. Low levels of viral RNA remained detectable in a subset of animals up to 365 dpi. Overall, visual inspection of the data did not reveal apparent differences in viral RNA kinetics among the Parental, Gamma, and Delta groups in any of the tissues analyzed, with substantial overlap in viral RNA levels across variants at all evaluated time points (Figure 4B).

Among the lymphoid organs, the thymus exhibited the highest viral RNA levels during the acute phase, reaching approximately 10³–10⁴ N2 copies/μL cDNA at 3 dpi. Viral RNA levels appeared to decline markedly by 15 dpi and remained detectable at low levels (approximately 10 N2 copies/μL cDNA) in a subset of animals up to 365 dpi (Figure 4C). In contrast, the spleen displayed the lowest viral RNA levels among all tissues analyzed. Viral RNA levels were already low at 3 dpi (approximately 10–100 N2 copies/μL cDNA) and remained within this range throughout the experimental period, without a clearly defined peak. Despite the low viral burden, viral RNA was still detectable in a subset of spleen samples at late time points, including 365 dpi (Figure 4D).

To further assess viral replication, genomic E and subgenomic E (sgE) RNA were also evaluated in the lungs, brain, spleen, and thymus. Both targets were detected exclusively during the acute phase of infection (3 dpi) in all analyzed tissues, whereas no amplification was observed at subsequent time points (Supplementary table 2). In contrast, N2 viral RNA remained detectable in a subset of animals up to 365 dpi, indicating prolonged persistence of viral RNA despite the absence of detectable E and sgE transcripts at later time points.

### SARS-CoV-2 nucleoprotein can persist in pulmonary and extrapulmonary tissues

The detection of viral RNA in pulmonary and extrapulmonary tissues at late time points led us to further investigate viral antigen persistence by immunofluorescence (IF). No specific nucleoprotein (NP) staining was observed in control (uninfected) animals in any of the tissues evaluated; in contrast, NP was detected in all infected groups, with the most intense immunostaining observed at 3 days post-infection (dpi). In the lung, intense staining was observed in the airway epithelium, along with focal signal in the thymus and spleen. Although NP-positive cells became progressively less frequent over time, staining persisted focally and at lower intensity in the lung and thymus at both 150 and 365 dpi, whereas spleen samples were not available for analysis at 365 dpi. Together, these findings corroborate the RT-qPCR data and further support the persistence of SARS-CoV-2 components in both pulmonary and extrapulmonary tissues well beyond the acute phase of infection (Figure 5).

**Figure 5.**
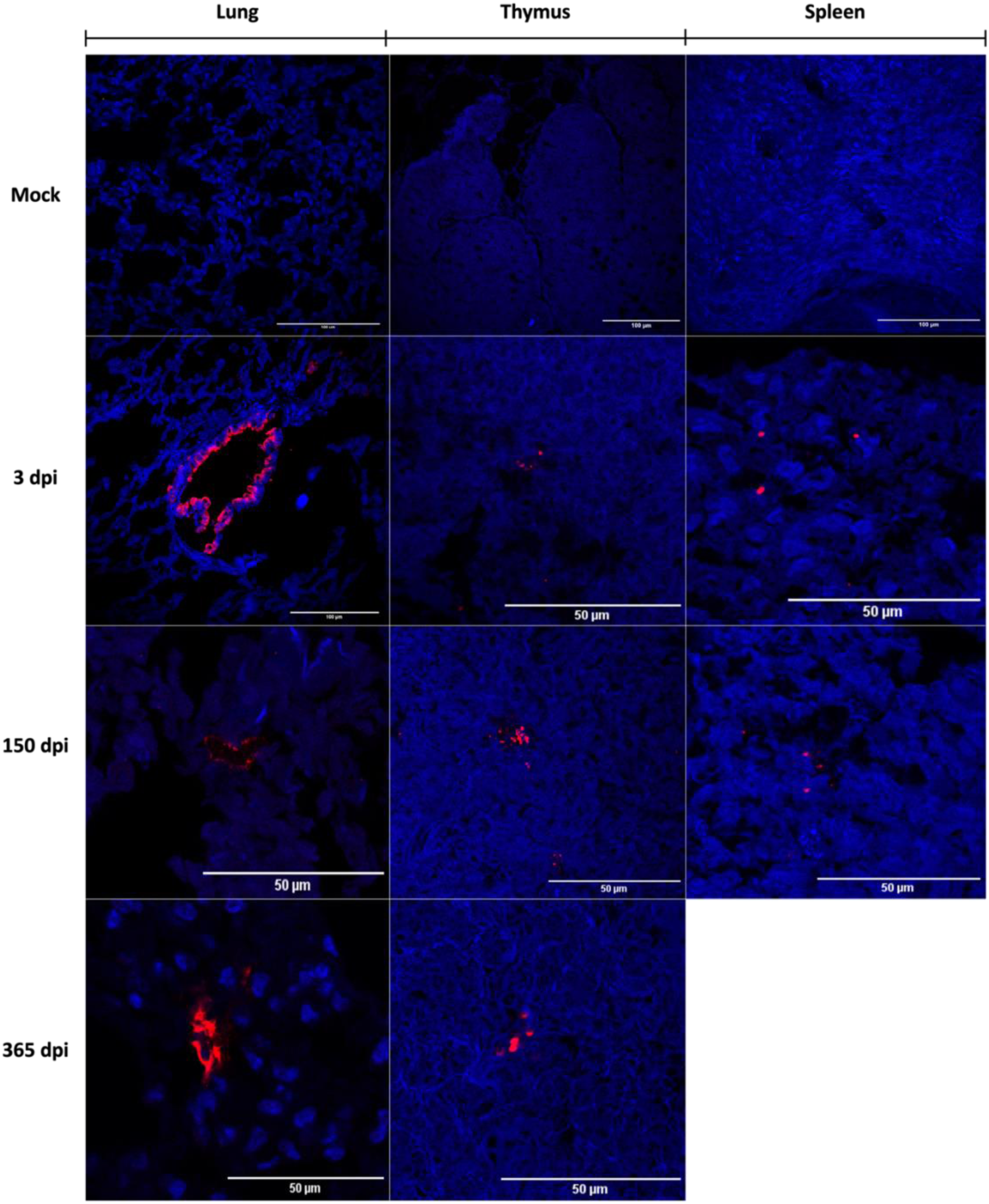
Detection of SARS-CoV-2 nucleoprotein by immunofluorescence in lung, thymus, and spleen throughout the course of infection. Representative immunofluorescence images showing viral nucleoprotein staining (red) and nuclear counterstaining with DAPI (blue) in lung, thymus, and spleen from mock-infected and infected golden hamsters at 3, 150, and 365 days post-infection (dpi), captured at 40× magnification. Scale bars: 100 μm (mock and lung at 3 dpi) and 50 μm (remaining panels).

### Temporal dynamics of cytokine mRNA expression across tissues following SARS-CoV-2 infection

The expression of immune-related genes was evaluated by RT-qPCR, and their relative expression in each tissue was normalized to the corresponding mock group at each time point and expressed as log₂ fold change.

The lung exhibited a well-defined temporal immune response, characterized by a robust acute inflammatory profile followed by a generalized suppression at later time points (Figure 6A). At 3 dpi, most immune-related genes were upregulated, with IL-2 showing the greatest induction among all targets analyzed, followed by CCL2, iNOS, TGF-β, and ISG15. Overall, the Delta variant displayed the strongest induction during the acute phase, particularly for IL-2, CCL2, and TNF-α, whereas CCL5, CXCL10, and IFN-γ showed only modest induction across all variant groups. By 150 dpi, the Parental and Gamma groups still exhibited mild upregulation of several immune-related genes, whereas the Delta group showed a trend toward reduced expression across multiple targets. At 365 dpi, a marked and widespread downregulation of nearly all immune-related genes was observed in the Parental, Gamma, and Delta groups relative to their respective mock controls, indicating long-term suppression of pulmonary immune gene expression. Among the analyzed targets, TGF-β appeared to be less affected than the other genes, although its expression remained below that of the mock group.

**Figure 6.**
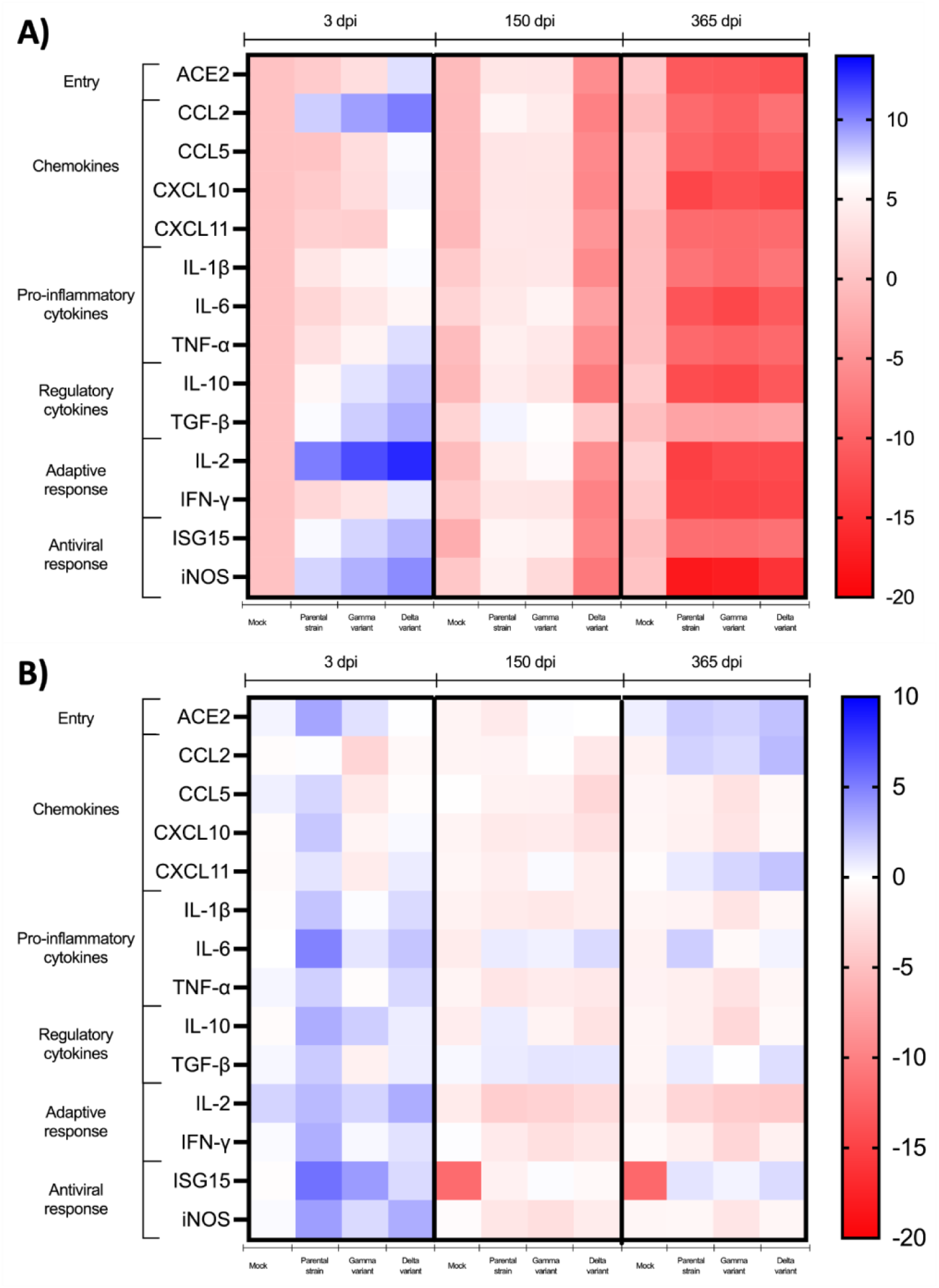
Heatmap of immune-related gene expression in the lung A) and brain B) of SARS-CoV-2-infected hamsters. Relative gene expression is shown as log₂(fold change) compared with the corresponding mock-infected group at 3, 150, and 365 dpi. Genes are grouped into viral entry factors, chemokines, pro-inflammatory cytokines, regulatory cytokines, adaptive immune response, and antiviral response. The color scale ranges from −20 (red, downregulation) to +14 (blue, upregulation) for the lung and from −20 to +10 for the brain, with white representing no change in gene expression.

In the brain, a more discreet overall inflammatory response was observed, characterized by a distinct late-phase chemokine signature rather than a uniform suppression of the immune response over time (Figure 6B). At 3 dpi, the magnitude of upregulation was lower compared to that observed in the lung in the parental strain-infected group, with the most notable increases observed for IL-6 and ISG15. Most chemokines and pro-inflammatory cytokines showed only mild changes relative to control animals. By 150 dpi, gene expression predominantly returned to near-baseline levels across all variant-infected groups, with only minor fluctuations observed. Notably, ISG15 expression was increased relative to mock-infected animals, in which ISG15 expression was undetectable at 150 and 365 dpi, consistent with its expected low basal expression in the absence of interferon stimulation. At 365 dpi, a distinct late immune activation signature emerged in all infected groups, characterized primarily by upregulation of CCL2, CXCL11, and ACE2 (Figure 6B).

In the thymus, immune-related gene expression followed a distinct temporal pattern, characterized by a delayed and pronounced activation peaking at 150 days post-infection rather than during the acute phase, followed by partial resolution at 365 dpi (Figure 7). At 3 dpi, changes in gene expression were generally modest across all variant-infected groups. Notably, the parental strain-infected group showed a slight reduction in CCL2 and IL-6 expression, while IFN-γ expression was moderately increased in this same group; most other targets remained close to levels observed in the mock control group. At 150 dpi, a marked and widespread increase in gene expression was observed across nearly the entire panel analyzed, particularly in the Delta variant-infected group, which showed the strongest induction among all groups for CXCL10, IL-1β, TNF-α, IL-2, IFN-γ, and iNOS. At 365 dpi, gene expression tended to return to baseline levels across all groups, although some alterations persisted, including downregulation of ACE2 and CCL5 in infected animals, increased IL-2 expression in the Delta group, and downregulation of ISG15 and TGF-β in infected groups (Figure 7)

**Figure 7.**
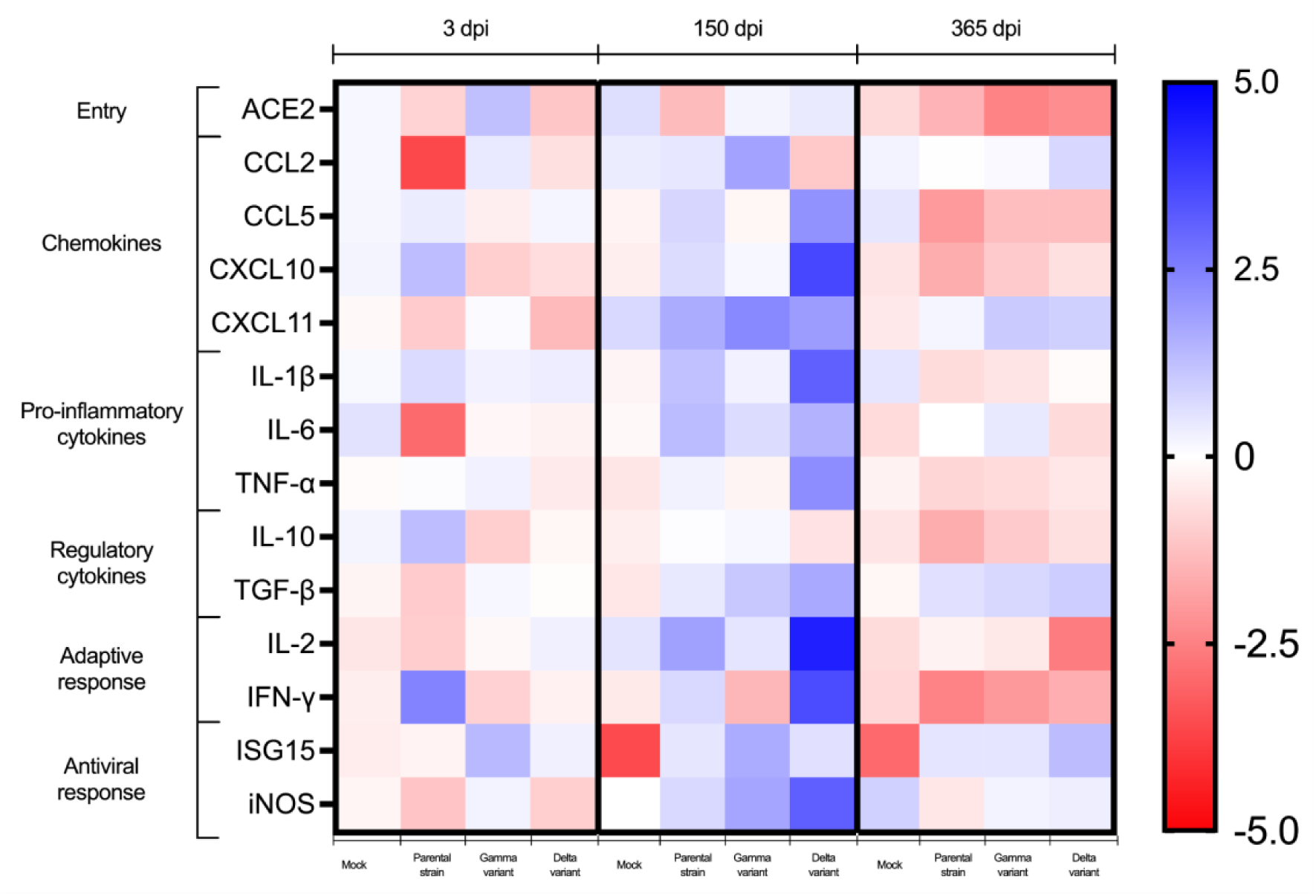
Heatmap of immune-related gene expression in the thymus of SARS-CoV-2-infected hamsters. Relative gene expression is shown as log₂(fold change) compared with the corresponding mock-infected group at 3, 150, and 365 dpi. Genes are grouped into viral entry factors, chemokines, pro-inflammatory cytokines, regulatory cytokines, adaptive immune response, and antiviral response. The color scale ranges from −5 (red, downregulation) to +5 (blue, upregulation), with white representing no change in gene expression.

### Sustained anti–SARS-CoV-2 IgG responses but limited cross-neutralization against the Omicron variant

To evaluate the humoral immune response induced by infection with different SARS-CoV-2 variants, anti–SARS-CoV-2 IgG levels were measured longitudinally up to 365 days post-infection. Infected animals exhibited detectable IgG responses from 15 days post infection (dpi), reaching peak concentrations at 30 dpi, with mean values ranging from approximately 1,200 to 1,600 ng/mL across the parental, Gamma, and Delta groups. Following the peak, a decrease in IgG levels was observed at 150 dpi. At 365 dpi, animals infected with the Gamma and Delta variants maintained higher IgG concentrations (approximately 1,400–1,600 ng/mL) compared to the parental strain (approximately 900–1,000 ng/mL). Mock-infected animals showed no detectable anti–SARS-CoV-2 IgG at any time point (Figure 8A).

**Figure 8.**
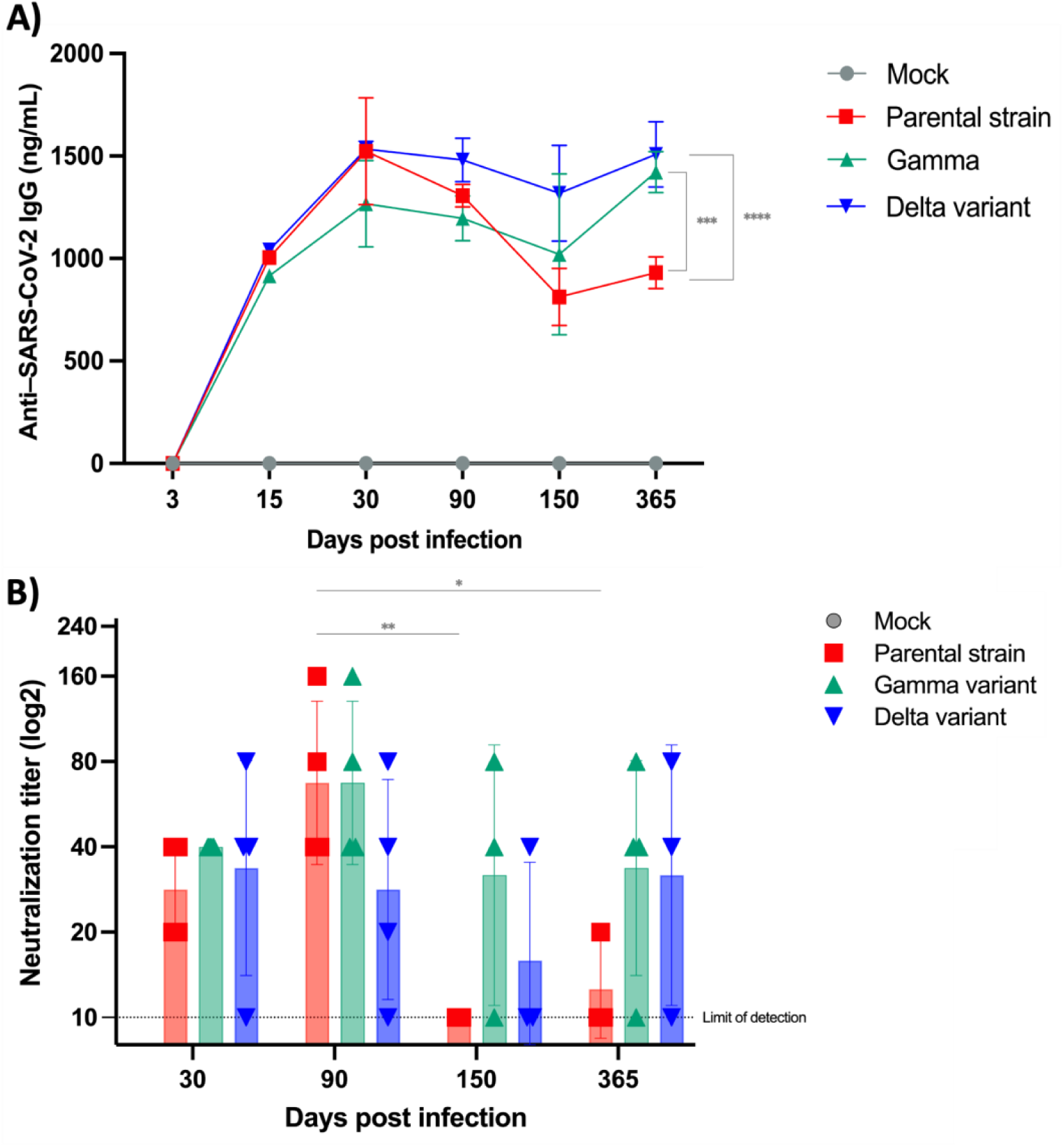
Longitudinal anti–SARS-CoV-2 humoral immune responses and cross-neutralization against the Omicron variant. **A)** Kinetics of anti-receptor binding domain (RBD) IgG antibody levels measured over 365 days post-infection. Antibody concentrations are expressed as ng/mL. “***p < 0.001, Parental strain vs. Gamma variant; ****p < 0.0001, Parental strain vs. Delta variant (365 dpi; two-way ANOVA with Tukey’s multiple comparisons test).“**B)** Serum microneutralization titers against the SARS-CoV-2 Omicron variant, demonstrating the development and persistence of cross-neutralizing antibody responses throughout the study period.

To evaluate cross-protective immunity against emerging SARS-CoV-2 variants, neutralizing antibody responses against the Omicron variant were assessed over time. Neutralizing antibodies against the Omicron variant were not detected in mock animals at any evaluated time point. In animals infected with pre-Omicron SARS-CoV-2 variants (Parental strain, Gamma, or Delta), neutralizing activity against Omicron emerged at approximately 30 days post-infection. The highest neutralizing titers were observed around 90 days post-infection, followed by a progressive decline over time. Despite this reduction, low levels of neutralizing activity persisted up to 365 days post-infection (Figure 8B). Overall, cross-neutralization titers remained relatively low, indicating limited cross-protective immunity against Omicron following infection with earlier SARS-CoV-2 variants. These findings are consistent with the well-described immune escape properties of the Omicron variant.

### SARS-CoV-2 infection induces long-term alterations in the serum metabolome, with a distinct signature associated with the Delta variant

Untargeted serum metabolomic analysis identified distinct metabolic signatures among animals infected with the Parental, Gamma, and Delta SARS-CoV-2 variants at both 3 and 365 dpi (Figure 9). Hierarchical clustering of significantly altered metabolites demonstrated clear differences in metabolite abundance patterns according to both viral variant and time post-infection.

**Figure 9.**
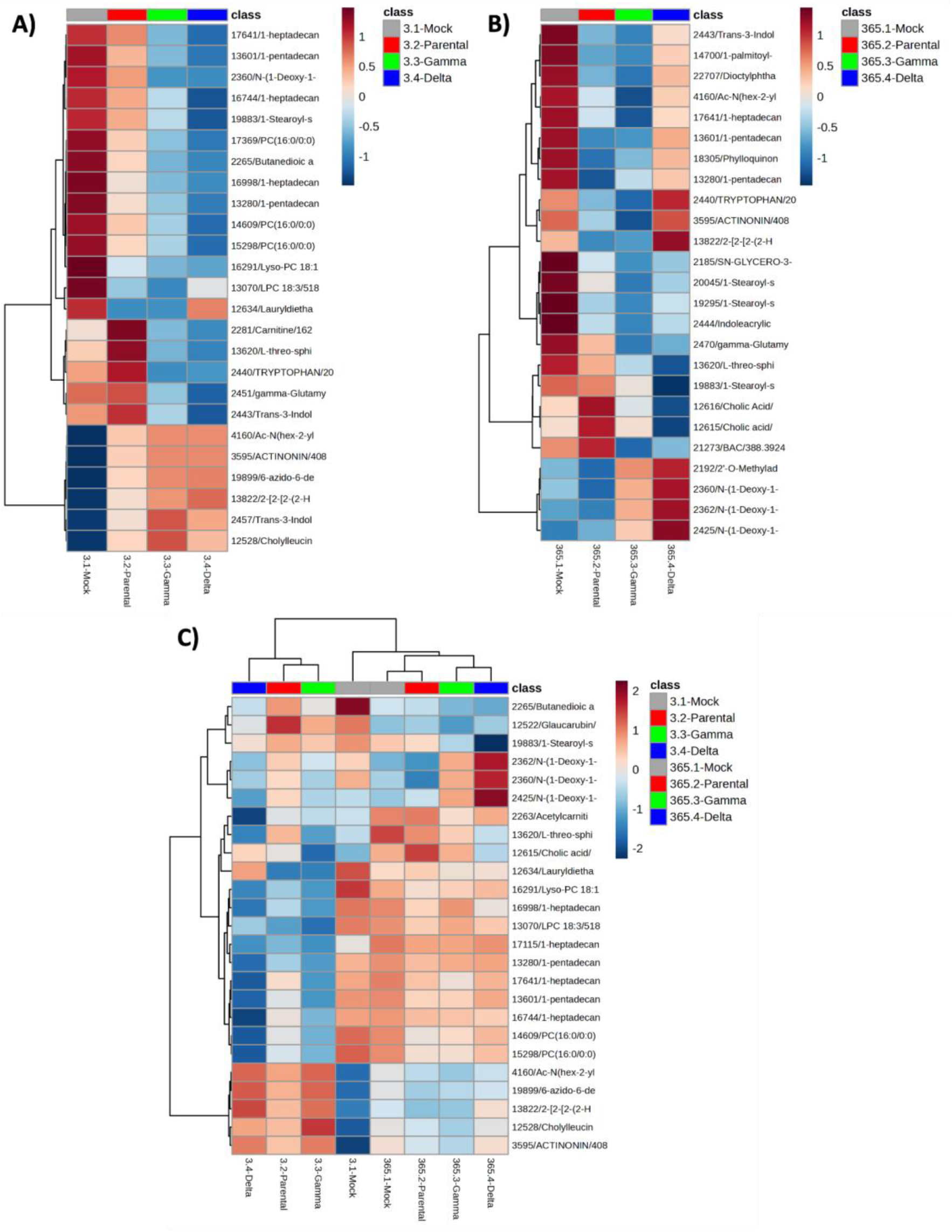
Global metabolomic profiles during acute and late SARS-CoV-2 infection. Hierarchical heatmaps showing differential metabolite abundance in serum from mock-, parental strain-, Gamma-, and Delta-infected hamsters. Data are presented as normalized values, with the color scale indicating relative metabolite abundance (red, higher; blue, lower). **A)** 3 days post-infection (dpi). **B)** 365 dpi. **C)** Combined analysis of samples from 3 and 365 dpi, highlighting metabolic changes between the acute and late phases of infection.

At 3 dpi, a clear separation was observed between the infected and control groups. Animals infected with the Gamma and Delta variants exhibited an overall reduction in the relative abundance of several metabolites compared to the Mock and Parental groups. Among the most notably altered compounds were lipid metabolism-related metabolites, including phosphatidylcholines (PCs), lysophosphatidylcholines (LPCs), and sphingolipid-associated species. Changes were also observed in metabolites associated with amino acid metabolism, particularly tryptophan-derived compounds and γ-glutamyl amino acid derivatives. Notably, the Parental group displayed a less pronounced metabolic perturbation relative to the variants of concern, suggesting a potential association between variant pathogenicity and the extent of host metabolic disruption (Figure 9A).

At 365 dpi, hierarchical clustering revealed variant-specific patterns of metabolic persistence rather than a uniform disruption across groups. Long-chain lipid species (palmitoyl- and pentadecanoic acid derivatives, phylloquinone) were reduced in both Gamma- and Delta-infected animals, with Gamma showing the most pronounced depletion. Tryptophan-derived metabolites, however, remained comparable to Mock in the Delta group, while Parental and Gamma showed sustained reductions. Phosphatidylcholine-related species and γ-glutamyl amino acid derivatives followed a graded reduction (Mock > Parental > Gamma ≈ Delta), suggesting a dose-dependent relationship with variant pathogenicity. Notably, sphingolipid and bile acid-related metabolites were selectively reduced in Delta alone, while a cluster of modified nucleosides showed the opposite pattern, with increased abundance exclusive to Delta (Figure 9B). Together, these findings indicate that Delta-infected animals display a distinct, and in some pathways divergent, metabolic signature at 365 dpi, suggesting that long-term metabolic disturbance is variant-specific rather than proportional to disease severity.

The integrated analysis of metabolic profiles at 3 and 365 dpi (Figure 9C) revealed that acutely infected samples formed a distinct cluster, clearly separated from all other groups, indicating a shared acute-phase metabolic signature across viral variants. In contrast, most groups evaluated at 365 dpi (Mock, Parental, and Gamma) clustered closely with the 3 dpi control group, suggesting a progressive resolution of the metabolic perturbations over time. However, the Delta group at 365 dpi formed a distinct subcluster, suggesting the persistence of residual metabolic alterations during the late phase of infection

## DISCUSSION

The in vitro infection of syrian hamsters proved, in this study, to be a good model for clarifying the pathogenesis mechanisms involved in SARS-CoV-2 infection and was already recommended in the literature due to its permissive system for clinical signs of the disease (Dai et al., 2025). When using different variants (Parental, Delta, and Gamma), changes in biomolecular signals can be observed depending on the inoculum. Animal body weight showed greater alteration following infection with the Parental and Gamma variants compared to the Delta variant, indicating that the Parental and Gamma variants are associated with more pronounced clinical effects regarding weight loss, whereas the Delta variant has a milder impact on this parameter.

The higher sensitivity of the N2 target is particularly relevant for the detection of low viral loads during late phases of infection and viral persistence in tissues, given its sensitivity in detecting viral RNA at a level of 0.2 TCID50. In our samples, genetic material was detected in all tissues during the acute phase and remained detectable, albeit at low levels, at later time points, consistent with the definition of viral persistence (Proal et al., 2025). The virus somehow manages to cross the blood-brain barrier and persistently infect brain tissue, as viral RNA and replicative virus have been detected in the brainstem 80 days after infection, generating unique transcriptomic profiles related to neurodegenerative processes (Coleon et al., 2025) In our cohort, the N2 target and the Envelope subgenomic RNA were also detected during the acute phase via RT-qPCR; however, only N2 remained detectable in brain tissue up to 365 days post-infection (dpi). Viral nucleoprotein was also found in all pulmonary and extrapulmonary tissues, showing a decline in frequency over time. These findings support the conclusion that the virus persists in various forms—whether as replication-competent particles, antigens, or replicating genetic material—corroborating mechanisms described in current literature regarding potential viral reservoirs in Long COVID (Prakash et al., 2025), which can lead to aftereffects that are still being documented by the scientific community and require further study (Lupi et al., 2024).

Animal serum was used to evaluate the humoral immune response, a high IgG response was observed starting fifteen days post-infection, followed by a drop in titers at 105 days and a recovery by the one-year mark. The Gamma and Delta variants maintained higher concentrations compared to the parental strain, indicating a sustained IgG humoral response. Regarding cross-neutralization with the Omicron variant, titers showed a progressive decline, consistent with the well-documented immune escape properties of the Omicron variant (Kudriavtsev et al., 2022).

Metabolomic profiling revealed extensive metabolic remodeling during the acute phase of SARS-CoV-2 infection, characterized primarily by alterations in lipid and amino acid metabolism. This metabolic signature, shared among the different viral variants and observed at 3 days post-infection (dpi), is consistent with previous reports describing common host metabolic responses to SARS-CoV-2 infection, including dysregulation of lipid and amino acid metabolism, as well as arachidonic acid, tryptophan, and their derived metabolites (Kaur, Ji & Rahman, 2021).

Although most metabolic alterations were reversed over time, Delta-infected animals maintained a distinct metabolic profile at 365 dpi, suggesting the presence of long-term variant-dependent metabolic effects. This finding resembles observations in individuals with post-COVID-19 condition, in which persistent alterations in phospholipid and lipid metabolism have been reported months or even years after acute infection, largely associated with changes in fatty acid metabolism and dysfunctional mitochondria-dependent lipid catabolism (López-Hernández et al., 2023; Yao et al., 2024).

This study has some limitations, including the restricted understanding of long-term SARS-CoV-2 persistence in the golden Syrian hamster model and differences between hamster and human immune responses that may limit direct extrapolation to post-acute COVID-19 sequelae. Additionally, the absence of consistent assessment of replication-competent viruses prevents definitive conclusions regarding active viral replication versus residual viral material. The evaluation of a limited number of variants may also not represent the full diversity of persistence patterns. Despite these limitations, the golden Syrian hamster remains a valuable model for studying long-term SARS-CoV-2 persistence and associated host responses. Future studies including additional variants, extended time points, and complementary approaches to evaluate viral replication and immune mechanisms will further clarify the role of viral persistence in post-acute COVID-19 outcomes.

In conclusion, SARS-CoV-2 infection in golden Syrian hamsters recapitulates key clinical and immunological features of COVID-19, including acute weight loss, robust pulmonary inflammation, and long-term persistence of viral RNA in multiple tissues for up to 365 days post-infection. Despite the absence of evidence for active viral replication, persistent viral RNA was associated with sustained immune and metabolic alterations. These findings highlight the value of the golden Syrian hamster as a model for investigating SARS-CoV-2 persistence and its potential contribution to the long-term consequences of COVID-19, including post-acute sequelae.

## ACKNOWLEDGMENTS

This work was supported by the São Paulo Research Foundation (FAPESP; grant no. 2019/26119-0), the Coordination for the Improvement of Higher Education Personnel (CAPES) (Finance Code 001), and the National Council for Scientific and Technological Development (CNPq). T.M.L. was supported by FAPESP (grant no. 2020/07063-1) and by the CAPES, (grant nos. 88887.480297/2020-00 and 88887.518483/2020-00).

We thank Jacqueline Nakau, specialist in charge of the Central de Espectrometria de Massas de Micromoléculas Orgânicas (CEMMO), School of Pharmaceutical Sciences of Ribeirão Preto, University of São Paulo (FCFRP-USP), for technical support with mass spectrometry analyses.

We also acknowledge Elizabete Rosa Milani, specialist in charge of the Laboratório Multiusuário de Microscopia Confocal (LMMC), supported by FAPESP (grant no. 2022/11494-3) for technical support with confocal microscopy.

We thank Soraya Jabur Badra, laboratory technician, for their technical support and assistance with biosafety level 3 (BSL-3) laboratory activities, including access to equipment and reagents at the Center for Research in Virology.

We are also grateful to Dr. Edison Durigon (Institute of Biomedical Sciences, University of São Paulo, São Paulo, Brazil) for providing the SARS-CoV-2 Gamma and Delta variants.

